# Dissociation between CSD-evoked metabolic perturbations and meningeal afferent activation and sensitization: implications for mechanisms of migraine headache onset

**DOI:** 10.1101/149278

**Authors:** Jun Zhao, Dan Levy

## Abstract

The onset of the headache phase during attacks of migraine with aura, which occur in about 30% of migraineurs, is believed to involve cortical spreading depression (CSD) and the ensuing activation and sensitization of primary afferent neurons that innervate the intracranial meninges, and their related large vessels. The mechanism by which CSD enhances the activity and mechanosensitivity of meningeal afferents remains poorly understood, but may involve cortical metabolic perturbations. We employed extracellular single-unit recording of meningeal afferent activity and monitored changes in cortical blood flow and tissue partial pressure of oxygen (tpO_2_) in anesthetized male rats to test whether the prolonged cortical hypoperfusion and reduction in tissue oxygenation that occur in the wake of CSD contribute to meningeal nociception. Suppression of CSD-evoked cortical hypoperfusion with the cyclooxygenase inhibitor naproxen blocked the reduction in cortical tpO_2_, but had no effect on the activation of meningeal afferents. Naproxen, however, distinctly prevented CSD-induced afferent mechanical sensitization. Counteracting the CSD-evoked persistent hypoperfusion and reduced tpO_2_ by preemptively increasing cortical blood flow using the K(ATP) channel opener levcromakalim did not inhibit the sensitization of meningeal afferents, but prevented their activation. Our data show that the cortical hypoperfusion and reduction in tpO_2_ that occur in the wake of CSD can be dissociated from the activation and mechanical sensitization of meningeal afferent responses suggesting that the metabolic changes do not contribute directly to these neuronal nociceptive responses.

**Significance statement:** CSD-evoked activation and mechanical sensitization of meningeal afferents is thought to mediate the headache phase in migraine with aura. We report that blocking the CSD-evoked cortical hypoperfusion and reduced tpO_2_ by cyclooxygenase inhibition is associated with the inhibition of the afferent sensitization but not their activation. Normalization of these CSD-evoked metabolic perturbations by activating K(ATP) channels is, however, associated with the inhibition of afferent activation but not sensitization. These results question the contribution of cortical metabolic perturbations to the triggering mechanism underlying meningeal nociception and the ensuing headache in migraine with aura, further point to distinct mechanisms underlying the activation and sensitization of meningeal afferents in migraine and highlight the need to target both processes for an effective migraine therapy.

## Introduction

Migraine is a multifactorial, episodic neurological disorder and the second leading cause of disability worldwide (Vos et al., 2017). It is now well accepted that the onset of migraine headache requires the activation and increased mechanosensitivity of trigeminal afferent neurons that innervate the intracranial meninges and their related large blood vessels (Strassman et al., 1996; Messlinger, 2009; Olesen et al., 2009; Levy, 2012). However, the endogenous processes underlying these afferent responses during a migraine attack are incompletely understood.

Migraine with aura, the second-most prevalent migraine subtype, is thought to involve cortical spreading depression (CSD), an abnormal wave of neuronal and glial activity (Ayata and Lauritzen, 2015). CSD has been suggested to mediate the aura phase (Pietrobon and Moskowitz, 2012), and also contributes to the onset of the headache phase, by promoting prolonged activation and increased mechanosensitivity of meningeal afferents (Zhang et al., 2010a; Zhao and Levy, 2015, 2016) and the ensuing activation and sensitization of central trigeminal dorsal horn neurons that receive their input (Zhang et al., 2011; Melo-Carrillo et al., 2017). The precise mechanisms by which CSD promotes the activation and sensitization of meningeal afferents are unclear, although dural neurogenic inflammation (Bolay et al., 2002) (but see however (Zhao and Levy, 2018)) and Pannexin-1 related signaling have been hypothesized (Karatas et al., 2013).

Migraine with aura is associated with cortical hemodynamic changes, in particular a sustained reduction in cortical blood flow contemporaneous with the onset of the headache phase (Olesen et al., 1981; Olesen et al., 1982; Lauritzen, 1984; Hadjikhani et al., 2001; Hansen and Schankin, 2017). In animal studies, CSD has been shown to promote a similar prolonged cortical hypoperfusion (Fabricius and Lauritzen, 1993; Fabricius et al., 1995; Piilgaard and Lauritzen, 2009). CSD also leads to transient cortical hypoxia followed by prolonged and milder reduction in cortical tissue oxygen tension (tpO_2_) concomitant with the cortical hypoperfusion phase (Takano et al., 2007; Piilgaard and Lauritzen, 2009).

Reduced tissue oxygenation during acute exposure to high altitude can promote headache and trigger a migraine attack (Appenzeller, 1994; Broessner et al., 2016). Normobaric hypoxia has also been shown to trigger migraine in susceptible individuals (Schoonman et al., 2006; Arngrim et al., 2016). These clinical observations together with the notion that reduced blood flow and tissue oxygenation can lead to enhanced activity and responsiveness of primary afferent nociceptors that innervate the cornea, viscera, muscle, and skin (Haupt et al., 1983; Mense and Stahnke, 1983; Longhurst et al., 1991; MacIver and Tanelian, 1992; Hillery et al., 2011) prompted us to examine whether the cortical hypoperfusion and associated reduction in tpO_2_ that occur following CSD might contribute to the prolonged activation and sensitization of meningeal afferents. Recent work has shown that inhibition of cortical cyclooxygenase abrogates CSD-evoked cortical hypoperfusion (Gariepy et al., 2017). Here, we investigated whether cyclooxygenase inhibition might also ameliorate the CSD-evoked decreases in tpO_2_ and interfere with the associated meningeal afferent responses. Having identified such effects, we further tested whether a preemptive increase in cortical blood flow to counteract the CSD-evoked hypoperfusion and reduced tpO_2_, using the K(ATP) channel opener levcromakalim (Kleppisch and Nelson, 1995; Shin et al., 2003) might also inhibit the enhanced responses of meningeal afferents. Our data suggest that while cyclooxygenase inhibition, and K(ATP) channel opening can counteract the CSD-related cortical hypoperfusion and reduced tpO_2_, they exert distinct inhibitory effects on associated activation and sensitization of meningeal afferents, thus questioning the direct contribution of these cortical metabolic perturbations to meningeal nociception and the onset of headache in migraine with aura.

## Materials and Methods

### Animals, anesthesia and surgical preparation

The current study employed Sprague-Dawley rats (males, 250–350 g, Taconic). Rats were housed 3 per cage under a 12 h light/dark cycle, in a temperature- and humidity-controlled room in the animal care facility of the Beth Israel Deaconess Medical Center, Boston. Food and water were available *ad libitum.* All experimental procedures followed the “guide for Care and use of Laboratory Animal resources (NIH publication No. 85-23 revised 1996), were approved by the Animal Care and Use Committee of the Beth Israel Deaconess Medical Center, and were in compliance with the ARRIVE (*Animal* Research: Reporting of In Vivo Experiments) guidelines. Animals were deeply anesthetized with urethane (1.5 g/kg, ip) and mounted on a stereotaxic frame (Kopf Instruments). Core temperature was kept at 37.5-38°C using a homoeothermic control system. Animals were intubated and breathed spontaneously room air enriched with O_2_. Physiological parameters were collected throughout the experiments using PhysioSuite (Kent Scientific) and CapStar-100 (CWE). Data used in this report were obtained from animals exhibiting physiological levels of oxygen saturation (>95%), heart rate (350–450 bpm), and end-tidal CO_2_ (3.5–4.5%). A saline-cooled dental drill was used to perform 3 separate craniotomies (Figure 1A). One craniotomy was used to expose the left transverse sinus and the posterior part of the superior sagittal sinus, as well as the adjacent cranial dura, extending ∼2 mm rostral to the transverse sinus. Another small craniotomy (1 × 1 mm) was made in the calvaria over the right hemisphere, and was centered 2 mm caudal and 2 mm lateral to Bregma to allow insertion of the recording electrode. An additional small burr hole (0.5-mm diameter) was drilled to expose a small area of the frontal cortex to induce CSD (Zhao and Levy, 2015, 2016). The exposed dura was bathed with a modified synthetic interstitial fluid containing 135 mM NaCl, 5 mM KCl, 1 mM MgCl_2_, 5 mM CaCl_2_, 10 mM glucose, and 10 mM HEPES, pH 7.2. The femoral vein was cannulated to allow for intravenous administration.

**Figure 1.**
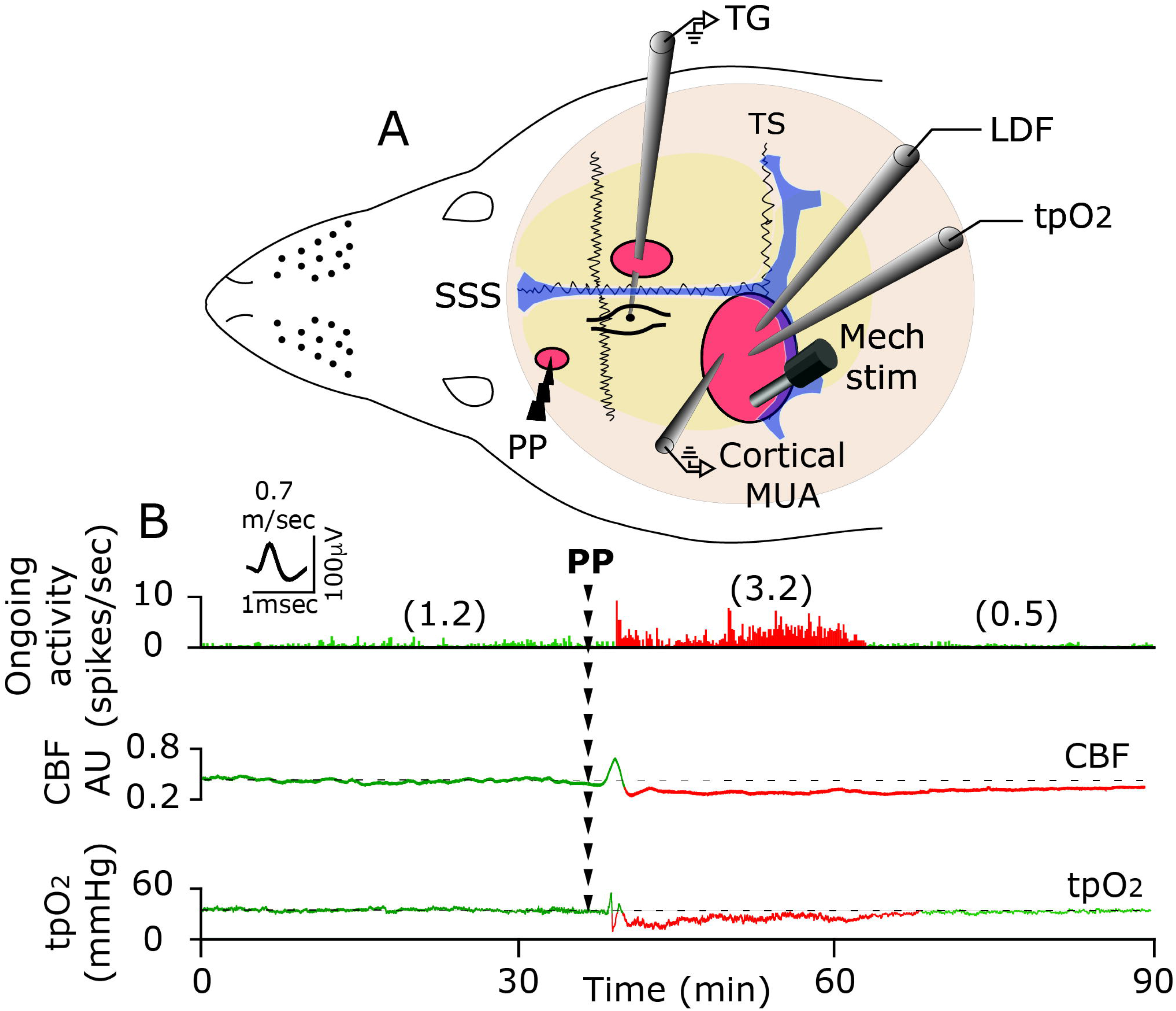
***A,*** Experimental setup: 3 skull openings (red ovals) were made. A small burr hole was made over the left frontal cortex to elicit a single cortical spreading depression (CSD) event using a pinprick (PP) stimulation. Meningeal afferent activity was recorded in the left trigeminal ganglion (TG) using a tungsten microelectrode inserted through a craniotomy made over the contralateral hemisphere. An ipsilateral craniotomy was made to expose a part of the left transverse sinus (TS) and superior sagittal sinus (SSS) and their vicinity to search for mechanosensitive meningeal afferents. Quantitative mechanical stimuli were delivered to the afferents receptive field using a feedback-controlled mechanical stimulator. Laser Doppler flowmetry (LDF) probe was placed over the cortex near the afferent’s receptive field to record changes in cortical blood flow and validate the induction of CSD non-invasively. In some animals, we also recorded the induction of CSD by monitoring multi-unit activity (MUA) of cortical neurons. An oxygen microelectrode was place in the superficial cortex to record CSD-evoked changes in cortical tissue partial pressure of oxygen (tpO_2_). ***B***, Example of raw data recording depicting the activation of a C-afferent meningeal afferent (with an overdrawn spike waveform used for data analysis) following the elicitation of CSD with a pinprick (PP) during the prolonged reduction in cortical blood flow (CBF) and tpO_2_. Afferent recording trace represents a peristimulus time histogram (PSTH, 1 second bin-size). Average ongoing activity rate (spikes/s) are denoted in parentheses. The activation phase and associated cortical hypoperfusion and reduced tpO_2_ are colored red. LDF values are in arbitrary units (AU).

### Recording of meningeal afferent activity

Single-unit activity of meningeal afferents (1 afferent/rat) was recorded from their cell body in the ipsilateral (left) trigeminal ganglion using a contralateral approach, as described recently (Zhao and Levy, 2014, 2015, 2016) (and see Figure 1A) in which a platinum-coated tungsten microelectrode (impedance 50-100 kΩ, FHC) is advanced through the right hemisphere into the left trigeminal ganglion using an angled (22.5°) trajectory. The insertion of the recording electrode using this approach does not produce CSD in the left cortical hemisphere (Zhao and Levy, 2015, 2016). Meningeal afferents were identified by their constant latency response to electrical stimuli applied to the dura above the ipsilateral transverse sinus (0.5-ms pulse, 5 mA, 0.5 Hz). The response latency was used to calculate conduction velocity (CV), based on a conduction distance to the trigeminal ganglion of 12.5 mm (Strassman et al., 1996). Neurons were classified as A-delta (1.5 ≤ CV ≤ 5 m/sec) or C afferents (CV < 1.5 m/sec). Neural activity was digitized and sampled at 10KHz using power 1401/spike 2 interface (CED). A real-time waveform discriminator (Spike 2 software, CED) was used to create and store a template for the action potential evoked by electrical stimulation, which was used to acquire and analyze afferent activity.

### Assessment of changes in meningeal afferent mechanosensitivity

Mechanical receptive fields of meningeal afferents were first identified by probing the exposed dura with a 2.0 g von Frey filament (Stoelting). The site of lowest mechanical threshold was further determined using monofilaments that exert lower forces (≥0.02 g). For quantitative determination of mechanical responsiveness, we applied mechanical stimuli to the site with the lowest threshold, using a servo force-controlled mechanical stimulator (Series 300B Dual Mode Servo System, Aurora Scientific), controlled via the power 1401 interface. The stimulus was delivered using a flat-ended cylindrical plastic probe that was attached to the tip of the stimulator arm. One of three probe diameters (0.5, 0.8 or 1.1 mm) was selected for each neuron, depending on the sensitivity of the neuron (Levy and Strassman, 2002). Stimulus trials for testing the afferent’s mechanosensitivity were made using a custom-written script for Spike 2 and consisted of ‘ramp and hold’ stimuli (rise time 100 msec, stimulus width 2 sec, inter-stimulus interval 120 sec). Each trial included a threshold stimulus (which normally evoked at baseline 1-3 Hz responses) followed by a supra-threshold stimulus (usually X2 of the threshold pressure). To minimize response desensitization, stimulus trials were delivered every 15 min throughout the experiment (Zhang et al., 2013; Zhao and Levy, 2016). Ongoing afferent discharge was recorded continuously between the stimulation trials. Responses to mechanical stimuli were determined during at least four consecutive trials before the elicitation of CSD. Data was analyzed only from afferents that exhibited consistent responses (variation of <0.5 Hz for threshold responses and <1.5 Hz for suprathreshold responses) during baseline recordings (Zhao and Levy, 2016).

### Induction of CSD

A single CSD episode was induced by quickly inserting a glass micropipette (diameter 50 μm) ∼2 mm deep in the cortex for 2 sec (Zhang et al., 2010a). CSD was triggered in the left frontal cortex to avoid potential damage to the meningeal tissue near the tested receptive field of the afferent under study, which could have led to their activation and/or sensitization (Zhao and Levy, 2015).

### Measurement of CSD-evoked changes in cortical blood flow using laser Doppler flowmetry

Changes in cortical blood flow (Gariepy et al., 2017) were documented using a needle probe (0.8 mm diameter) connected to a laser Doppler flowmeter (Vasamedic, St Paul MN). The probe was positioned just above the intact dura, ∼1 mm from the mechanical receptive field of the recorded afferent in an area devoid of large pial blood vessels (>100 μm). Such system (using a near IR 785 nm probe and fiber separation of 0.25 mm) is expected to record blood flow changes at a cortical depth of ∼1 mm (Fabricius et al., 1997). The laser Doppler signal was digitized (100Hz) and recorded using power 1401/Spike 2. Laser Doppler recordings were obtained with the ambient lights turned off. Baseline data were based on recording conducted for at least 30 min prior to CSD.

### Measurement of cortical partial pressure of oxygen (tpO_2_)

Cortical tissue partial pressure of O**_2_** (tpO_2_) was recorded using a modified Clark-type polarographic oxygen microelectrode (OX-10; Unisense) with a guard cathode. The oxygen electrode was connected to a high-impedance Pico-amperometer (PA2000; Unisense) that sensed the currents of the oxygen electrodes. The tpO_2_ signal was then digitized (100 Hz) and recorded using the Power 1401/Spike 2 interface. The microelectrode responded linearly to changes in oxygen concentration and was calibrated before and after each experiment in both air-saturated saline and in oxygen-free solution consisting of 0.1 M of sodium-ascorbate and 0.1 M NaOH dissolved in saline. This approach has been used previously to record changes in cortical tpO_2_ in response to CSD (Takano et al., 2007; Piilgaard and Lauritzen, 2009; Thrane et al., 2013). Changes in tpO**_2_** values were recorded at a cortical depth of 100-200 μm, with the electrode placed <0.5 mm away from the LDF probe. Baseline tpO_2_ values were recorded for at least 30 min prior to CSD.

### Electrophysiological recording of CSD

A tungsten microelectrode (impedance 1-2Ω, FHC), was used to record multi-unit activity at a depth of 200-400 μM in the left visual cortex (S1). A silver ground electrode was placed in the neck muscle. Signal was amplified and filtered (X20K, 300Hz low frequency; 10KHz high frequency, DAM-80/BMA400, WPI/CWE), and then digitized and sampled at 10KHz. Cortical activity was recorded simultaneously with CBF changes, with the recording electrode placed <0.5 mm away from the LDF probe.

### Drugs

Naproxen sodium and Levcromakalim were purchased from Tocris Biosciences. Naproxen sodium was made in 0.9% immediately before administration (10 mg/kg i.v). Levcromakalim stock solution (10 mM) was made in DMSO and further diluted with synthetic interstitial fluid (final concentration 500μM) immediately before its application. The concentration of levcromakalim expected to be attained at the superficial cortex was estimated to be 100-1000 times lower, (Moore et al., 1982; Roth et al., 2014; Zhao et al., 2016).

### Data analyses and statistics

All data were analyzed and plotted using Prism 7 software and is presented as [median (IQR)]. Group data in graphs show all data points and are plotted as box and whiskers with min and max values. For CSD-related changes in cortical blood flow and tpO_2_, raw data were processed with a custom-written script made for Spike 2. Baseline values were based on the average of the values collected during the 600 secs prior to the induction of CSD. Offline analyses for afferent responses were conducted using template matching in Spike 2. Criteria used to consider meningeal afferent activation and sensitization responses were based on our previous studies (Zhao and Levy, 2015, 2016). In brief, prolonged increase in afferents’ ongoing activity was considered if the firing rate increased above the upper end point of the 95% CI calculated for the baseline mean for >10 min. Acute afferent activation during the CSD wave was considered as an increased ongoing discharge rate that occurred ≥30 sec following the pinprick and lasted ≤120 sec. Mechanical sensitization was considered only if the afferent responses changed according to the following criteria: threshold and/or suprathreshold responses increased to a level greater than the upper endpoint of the 95% CI calculated for the baseline mean; increased responsiveness began during the first 60 min post-CSD; and sensitization lasted for at least 30 min. Changes in CSD-evoked responses of meningeal afferents in the presence of naproxen or levcromakalim were compared to data obtained in control animals using the same surgical approach and CSD induction method. Control data was obtained from experiments in animals treated intravenously with the naproxen vehicle (0.9% saline, 60 min prior to CSD induction, n=24), or locally (i.e. continuous dural bathing) with synthetic interstitial fluid (n=36). The two data sets were comparable and were combined to increase power. Differences in CSD-evoked afferent activation and sensitization propensities were analyzed using two-tailed, chi-square tests. Within- and between group comparisons were made using Wilcoxon test and Mann Whitney U-test respectively. Results were considered to be significant at *p*<0.05.

## Results

### CSD-evoked prolonged activation of meningeal afferents coincides with cortical hypoperfusion and reduced tpO_2_

Protracted cortical hypoperfusion and reduced cortical tpO**_2_** were observed previously (Piilgaard and Lauritzen, 2009; Fordsmann et al., 2013), at time points corresponding to the prolonged activation of meningeal afferents (Zhang et al., 2010a; Zhao and Levy, 2015). To examine whether the CSD-evoked cortical metabolic perturbations indeed coincide with prolonged meningeal afferent activation, we employed simultaneous recording of meningeal afferent activity with cortical blood flow and tpO**_2_** (n=6). As the example in Figure 1B demonstrates, persistent meningeal afferent activation could be observed during the prolonged post-CSD cortical hypoperfusion and reduced tpO_2_ stage.

### Suppression of CSD-evoked cortical hypoperfusion and decreased tpO_2_ by naproxen is not associated with reduced activation of meningeal afferents

We initially verified that systemic administration of naproxen, which was employed to ensure cortical COX-inhibition also under the nerve endings of meningeal afferents branches localized to areas not exposed by the craniotomy, could also block the CSD-evoked cortical hypoperfusion, as we previously have shown using topical administration (Gariepy et al., 2017). Because cortical oxygenation level depends in large part on cortical perfusion (Offenhauser et al., 2005), we further examined whether blocking the CSD-evoked cortical hypoperfusion with naproxen might inhibit the reduction in cortical tpO**_2._** When compared to control animals (n=22), naproxen treatment (n=22) augmented the CSD-related acute cortical hyperemic response and blocked, as expected, the protracted cortical hypoperfusion (Figure 2). Using simultaneous recording of cortical blood flow and tpO_2_, we observed that the inhibitory effect of naproxen on the CSD-evoked cortical hypoperfusion was associated with altered tpO**_2_** responses to CSD (Figure 3). In the presence of naproxen (n=6), CSD evoked an initial increase in tpO_2_ [Δ10.03 mmHg (5.22)] [median (IQR)]; (W=6.0; p=0.03; phase 1 in Figure 3B, and arrowhead in the insert) that was smaller than in vehicle-control animals (n=11); [Δ14.18 mmHg (3.87)] [median (IQR)]; (W=66.0; p=0.006; U=7.0; p=0.007, Figure 3C). While in control animals, the hyperoxia phase was followed by acute hypoxia; bottom values [17.66 mmHg (18.64)] [median (IQR)]; (W= −66.0; p=0.001), which lasted [110.0 sec (70.30)] [median (IQR)]; (phase 2, Figure 3A), naproxen blunted this hypoxic response, and tpO_2_ values increased instead [113.4 mmHg (90.24)] [median (IQR)]; (W=21.0; p=0.015; phase 2, Figure 3B). This hyperoxia phase returned to baseline levels at 15-25 min [59.02 mmHg (39.63)] [median (IQR)]; (W=7.0; p=0.29; phase 3, Figure 3B). Overall, when compared to control animals, naproxen inhibited both the acute hypoxia (phase 2; U=0; p<0.001; Figure 3D) and prolonged decrease in tpO_2_ (phase 3; U=3; p=0.005; Figure 3E).

**Figure 2.**
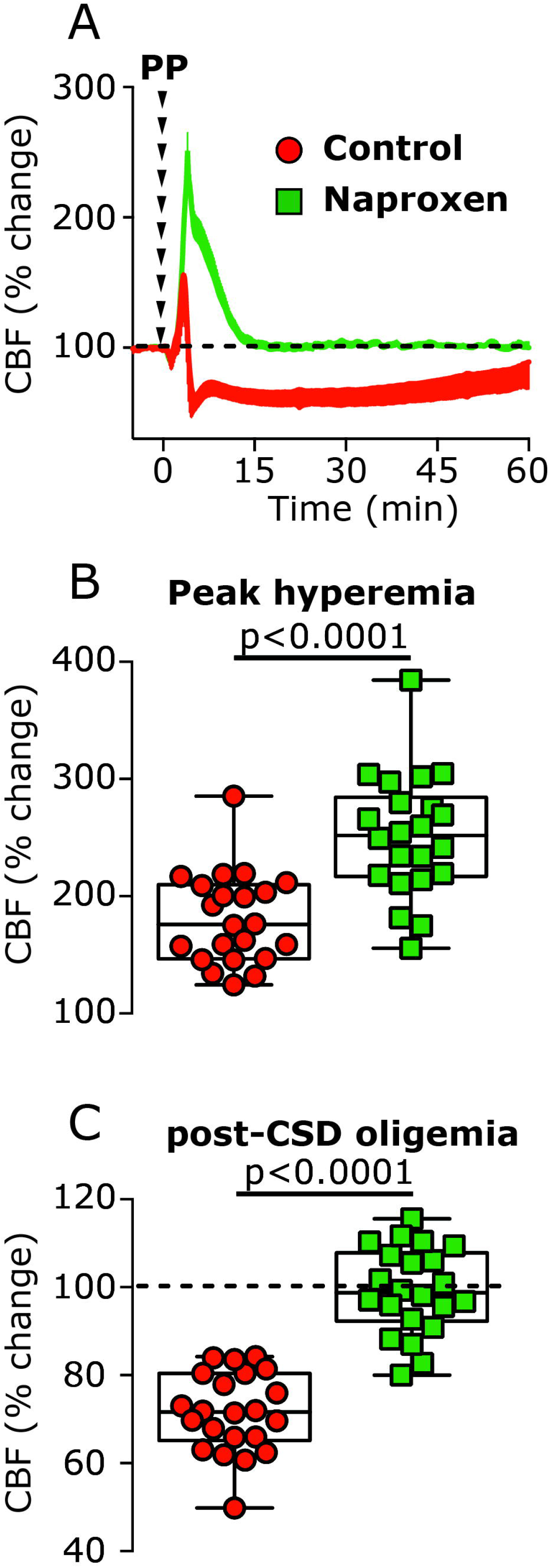
The effect of systemic naproxen on CSD-induced changes in cortical blood flow (CBF). ***A***, Changes in CBF following induction of CSD in vehicle- and naproxen-treated animals. **C**, CBF changes during the hyperemia stage, showing the enhancing effect of naproxen. **D,** CBF changes during the hypoperfusion phase, 10-60 min following CSD, depicting the inhibitory effect of naproxen. P values based on a Mann Whitney U-test.

**Figure 3.**
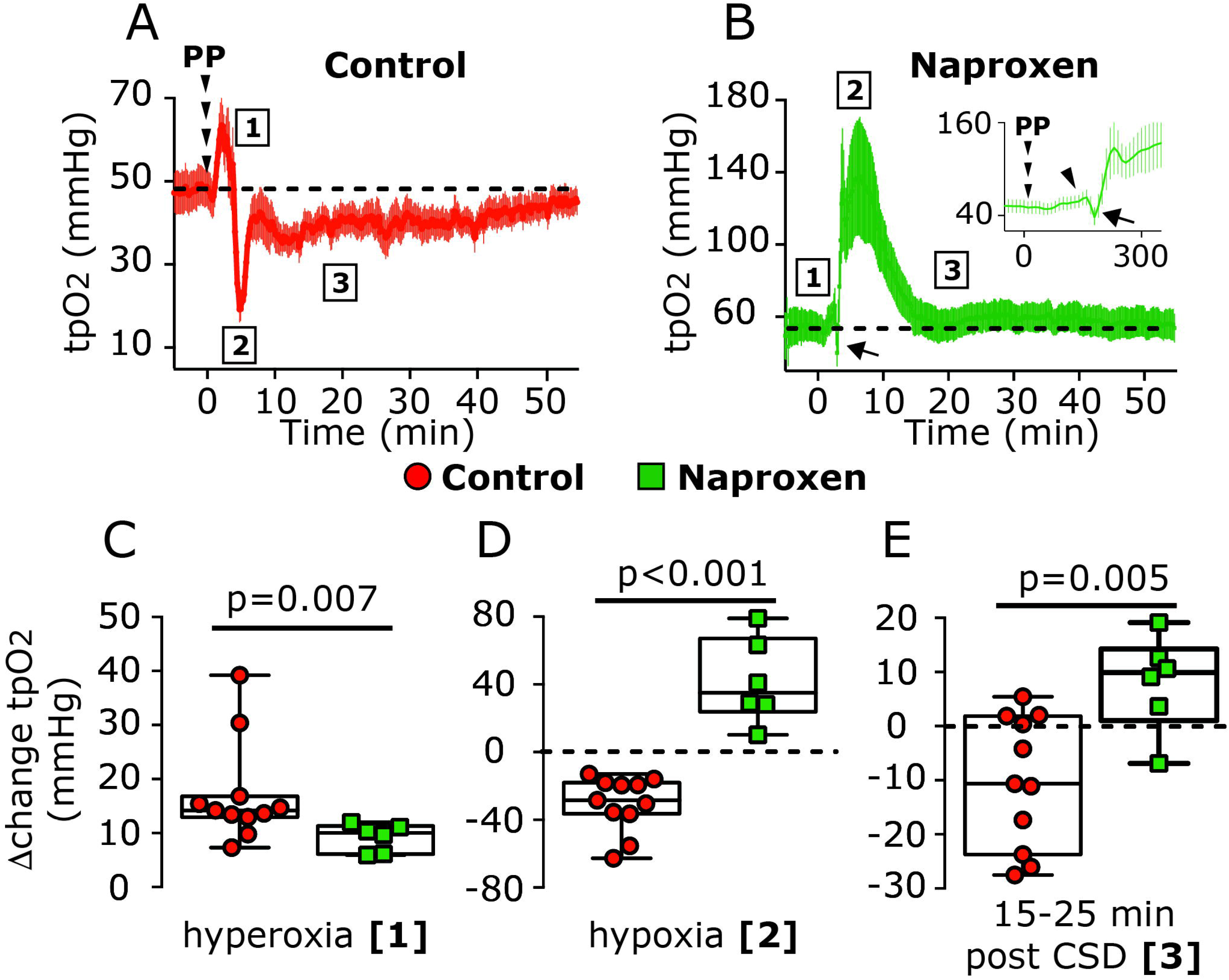
Naproxen inhibits CSD-induced decreases in cortical tpO_2._ Time– course of changes in tpO_2_ following CSD induction in control (***A***) and naproxen-treated animals (***B***). Comparisons between CSD-evoked changes in control and naproxen-treated animals during the hyperoxia phase (phase 1), the acute hypoxia phase (phase 2, in control animals) and at 15-25 min post-CSD (phase 3) (***C, D, E***). P values based on a Mann Whitney U-test.

We next examined whether the suppressing effects of naproxen on CSD-evoked cortical hypoperfusion and decreased tpO_2_ might be associated with reduced activation of meningeal afferents. A-delta afferents recorded from naproxen-treated animals (n=15) had similar baseline ongoing activity as in control, vehicle-treated animals (n=26); [0.04 spikes/sec (0.20)] vs [0.05 spikes/sec (0.25)] [median (IQR)]; (U=189.0; p=0.87). Baseline ongoing activity of C afferents were also not different between the naproxen (n=16) and control groups (n=34) [1.08 spikes/sec (1.58)] vs [0.57 spikes/sec (1.35)] [median (IQR)]; (U=215.0; p=0.24). In naproxen-treated animals, CSD evoked acute activation in 5/15 A-delta and 6/16 C, ratios which were not significantly different than those observed in control animals (9/26 A-delta, 14/34 C; χ^2^ =0.007; p=0.99 for the A-delta population; χ^2^ =0.06; p=0.80 for the C afferent populations; χ^2^=0.07; p=0.79 for the combined population). When compared to the control group, naproxen also did not affect the magnitude of the immediate activation, neither in A-delta [4.2 fold (3.50)] vs [3.6 fold (2.10)] [median (IQR)]; (U=22.0; p=0.99) nor in C afferents [1.6 fold (0.50)] vs [2.1 fold (1.40)] [median (IQR)]; (U=30.0; p=0.35, Figure 4C).

**Figure 4.**
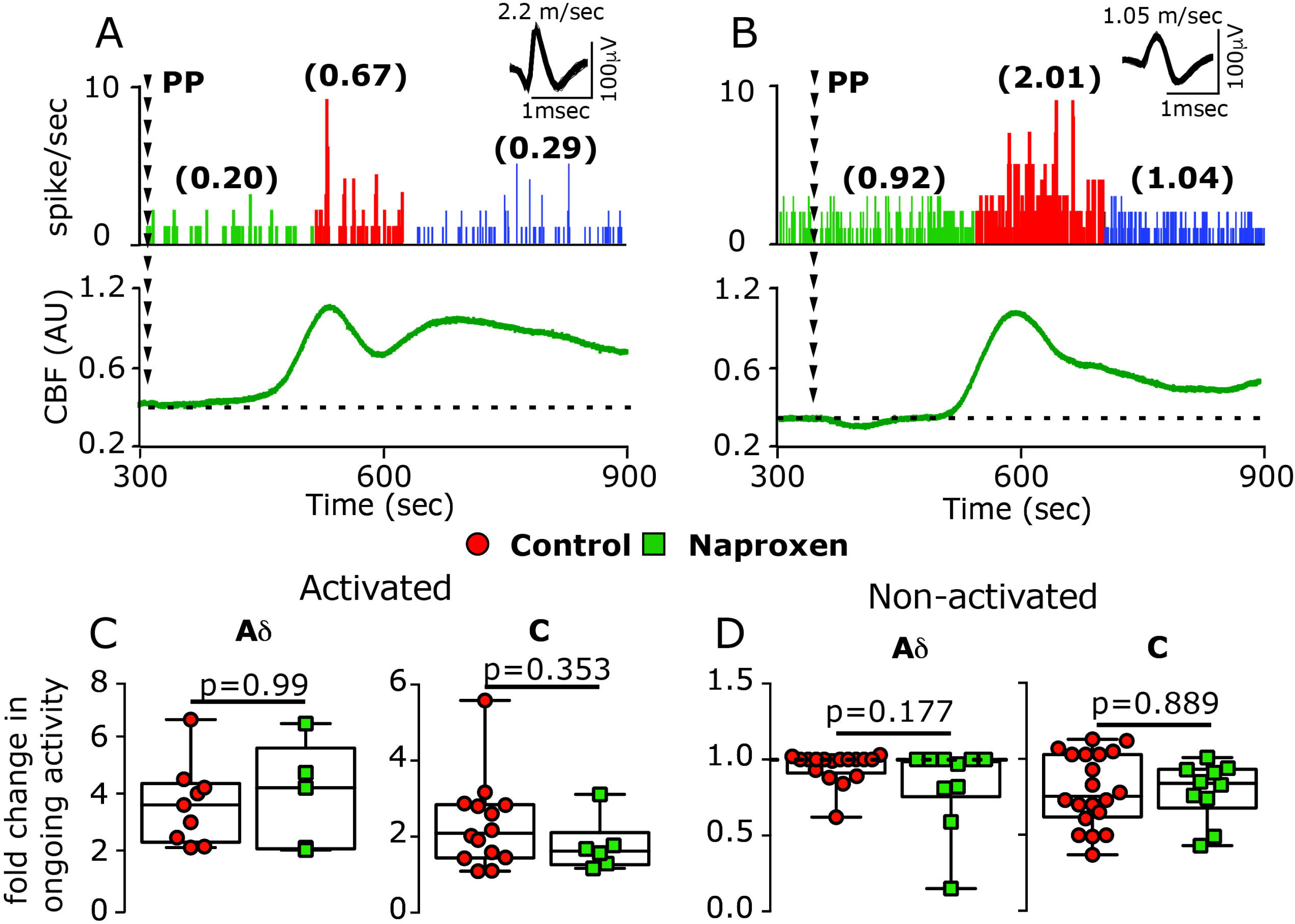
Naproxen treatment does not inhibit CSD-evoked acute activation of meningeal afferents. Examples of experimental trials depicting the acute activation of an A-delta afferent (***A***) and a C afferent (***B***) following the induction of CSD using a pinprick (PP) stimulus in animals treated with naproxen. Traces represent post-stimulus time histogram (1 second bin-size). Average ongoing activity rate (spikes/sec) is denoted in parentheses. Red areas represent the acute phase of afferent activation during CSD. Raw LDF traces are provided below each recording showing the CSD-evoked increase in CBF. Comparisons of the magnitude of change in ongoing activity between afferents recorded in control and naproxen-treated animals depicting activated (***C***) and non-activated afferents (***D***). P values based on a Mann Whitney U-test.

The likelihood of developing a prolonged increase in ongoing afferent activity following CSD in naproxen-treated animals (see examples in Figure 5A, B) was also not different when compared to that observed in control animals (7/15 vs 13/26, χ^2^ =0.04;2 p=0.84 for the A-delta; 7/16 vs 22/34, χ^2^ =1.96; p=0.16 for the C-afferents; χ^2^ =1.427; p=0.23 for the combined population). Naproxen treatment also did not affect any of the CSD-evoked afferent activation parameters: persistent activation developed in A-delta afferents with a delay of [25 min (40)] [median (IQR)], no different than in control animals [10 min (5.0)] [median (IQR)]; (U=33.0; p=0.34, Figure 5C); The activation delay observed in C afferents in naproxen-treated animals [15 min (31.25)] [median (IQR)] was also not different that in controls [5 min (17.5)] [median (IQR)]; (U=67.5; p=0.58, Figure 5C). The duration of CSD-evoked prolonged activation was also unaffected by naproxen: A-delta afferents were activated for 15 min (15.0)] vs [30 min (22.5)] [median (IQR)] in the control group (U=22.0; p=0.07, Figure 5D); C-afferents were activated for [30 min (31.25)] vs [35 min (30.0)] [median (IQR)] in the control group (U=58.0; p=0.34, Figure 5D). Finally, when compared to controls, naproxen treatment did not reduce the magnitude of the afferent activation observed in either population [A-delta; 2.3 fold (3.0)] vs [3 fold (3.8)] [median (IQR)]; (U=39.0; p=0.64); C afferents; [1.8 fold (3.0)] vs. [1.5 fold (0.9)] [median (IQR)]; (U=51.0; p=0.18, Figure 5E). In afferents deemed not-activated by CSD, the changes in ongoing activity following CSD were also not different between the naproxen and control groups, in both the A-delta (U=41.0; p=0.8) and C afferent (U=35.0; p=0.2) populations (Figure 5F).

**Figure 5.**
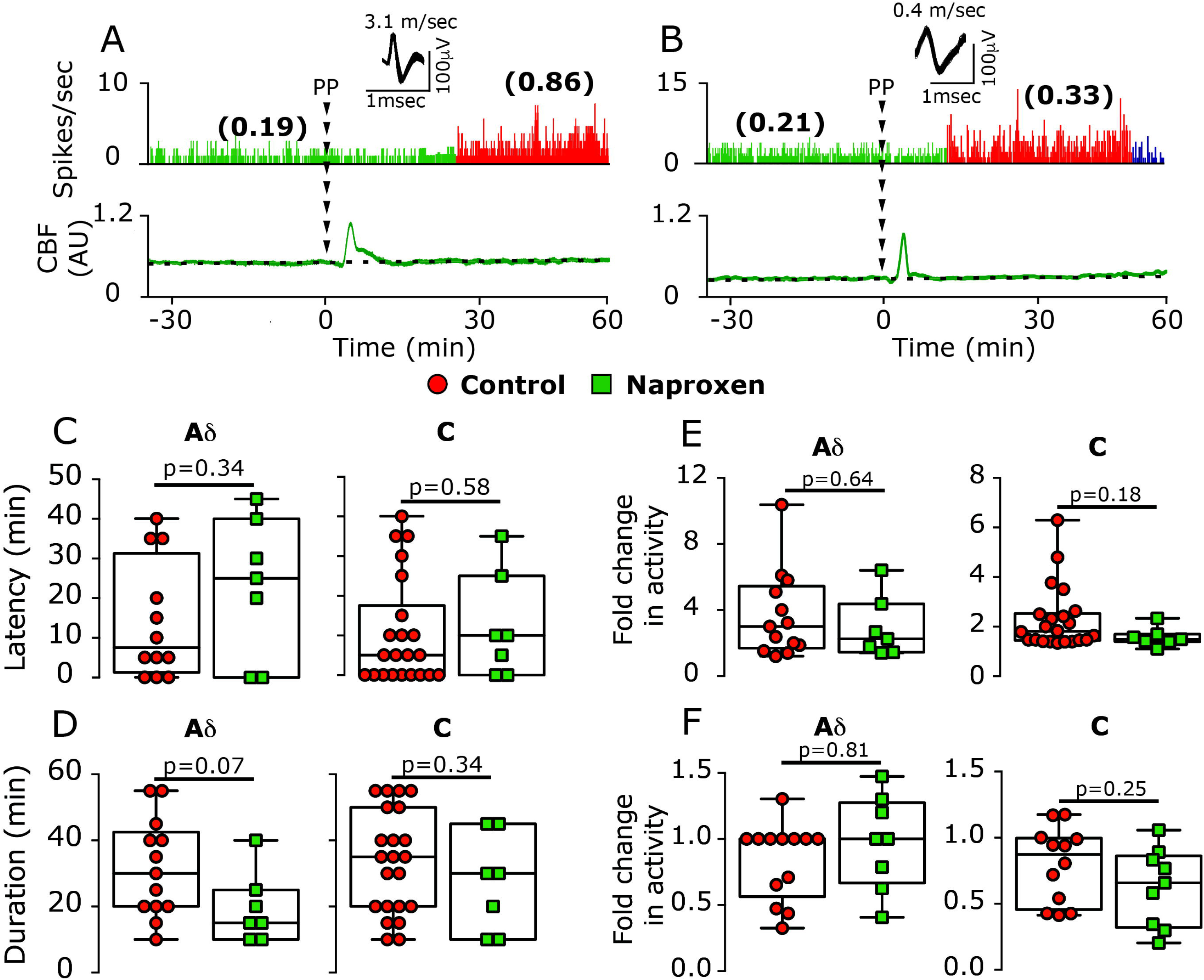
Naproxen treatment does not inhibit CSD-evoked prolonged activation of meningeal afferents. Experimental trials depicting the CSD-evoked prolonged activation of an A-delta afferent (***A***) and a C afferent (***B***) in animals treated with naproxen. Naproxen treatment did not affect the onset latency of the afferent activation (***C***), its duration (***D***), or magnitude (**E**). Changes in ongoing activity rate in non-activated afferents in naproxen-treated animals were not different than those in controls (***F***). P values based on a Mann Whitney U-test.

### Naproxen inhibits CSD-evoked meningeal afferent mechanosensitization

We previously proposed that the activation and mechanosensitization of meningeal afferents that occur following CSD are mediated via distinct mechanisms (Zhao and Levy, 2016). We therefore examined next in 8 A-delta and 7 C afferents whether the inhibitory effects of naproxen on the cortical hypoperfusion and decreased tpO_2_ might be associated with amelioration of the post-CSD mechanosensitization response. Because the characteristics of the CSD-evoked sensitizations of the A-delta and C afferents are similar (Zhao and Levy, 2016), we combined the data from the two afferent populations for comparisons with the data obtained from vehicle-treated animals (10 A-delta and 13 C). At baseline, the number of dural mechanosensitive receptive fields, and mechanical responsiveness of afferents recorded in naproxen-treated animals were similar to those in the control group (Table 1). Unlike its ability to inhibit the CSD-evoked activation of meningeal afferents, naproxen blocked the afferent sensitization responses, at both the threshold [(3/15 vs 13/23); χ(1) = 4.97; p=0.03; Figure 6C] and suprathreshold levels [(2/15 vs 13/23); χ(1) = 7.09; p=0.008; Figure 6]. When sensitization was defined as enhanced responsiveness at either stimulus level, there were also fewer sensitized afferents in the naproxen group [(4/15 vs 17/23); χ(1) = 9.084; p<0.003]. When the responses of all sensitized and non-sensitized units were combined, naproxen also had an overall inhibitory effect on the changes in threshold responses [Δ0 spike/sec (0.6)] vs [Δ0.39 spikes/sec (1.43)] [median (IQR)]; (U=90.5; p=0.01, Figure 6G) and suprathreshold responses [0.86 fold (0.5)] vs [1.2 fold (0.5)] [median (IQR)]; (U=22.0; p=0.004, Figure 6H).

**Table 1:** Baseline mechanical response properties of meningeal afferents in animals treated with vehicle, naproxen, or levcromakalim.

|  | **Receptive**<br>**fields <sup>a</sup>** | **Threshold**<br>**responses <sup>b</sup>** | **Suprathreshold**<br>**responses <sup>c</sup>** |
| --- | --- | --- | --- |
| **Control** | [2 (1)] | [1.3 (0.7)] | [7.3 (7.0)] |
| **Naproxen** | [2 (1)] | [0.5 (1.8)] | [8.0 (6.5)] |
| **Levcromakalim** | [2 (1)] | [1.7 (1.5)] | [9.0 (4.0)] |

Data are expressed as [**median** (IQR)]. Threshold and suprathreshold neural responses (spikes/sec) are based on activity recorded during a 2-second mechanical stimulus, and represent the average of 3-4 trials conducted prior to the elicitation of CSD in the presence of the vehicle, naproxen or levcromakalim. Group differences were analyzed using Kruskal-Wallis and were not statistically significant for the 3 parameters examined. **a** $H(2)=3.85$ , $p=0.145$ ; **b** $H(2)=4.70$ , $p=0.095$ ; **c** $H(2)=3.24$ , $p=0.198$ .

**Figure 6.**
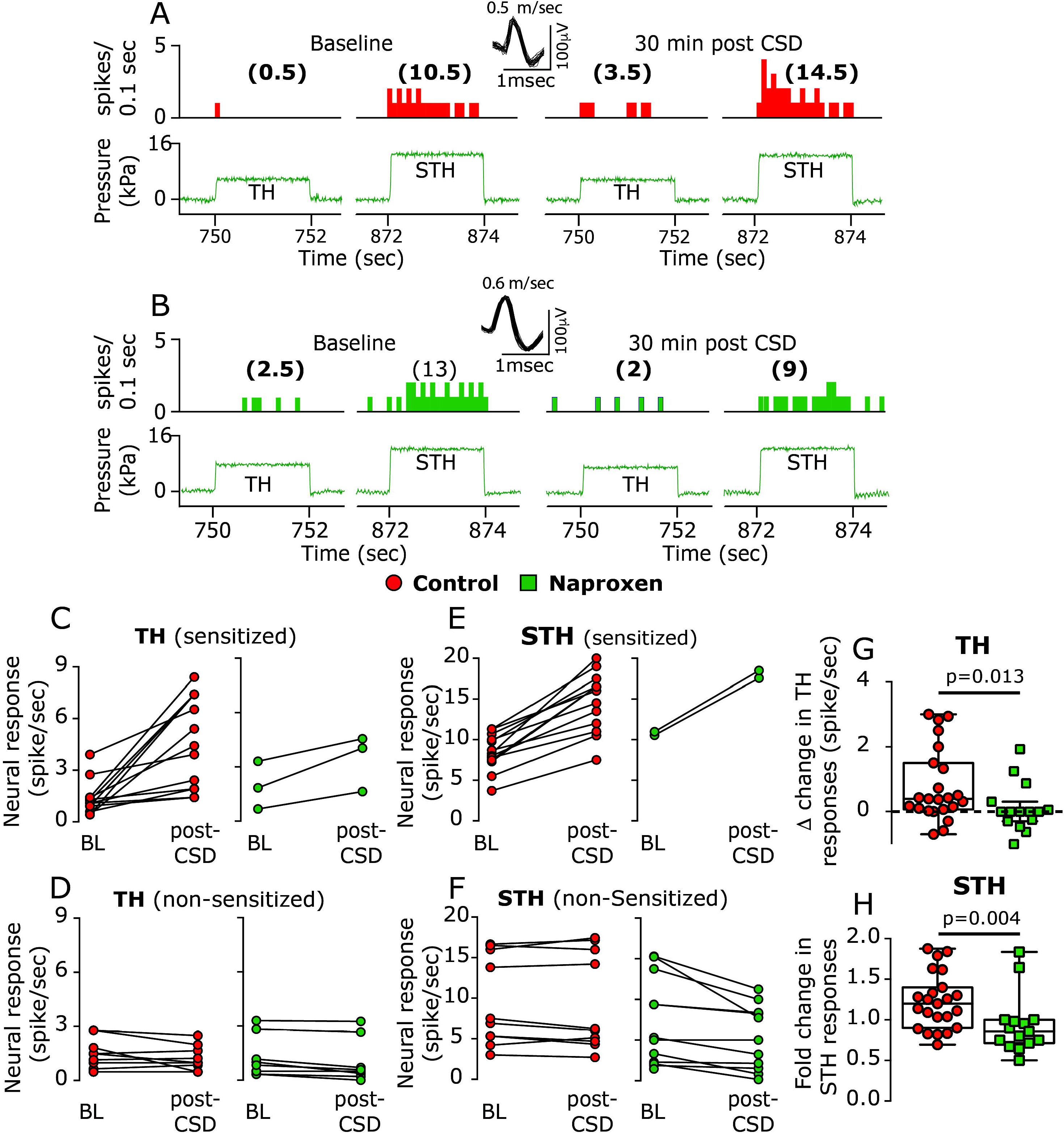
Naproxen inhibits CSD-induced mechanical sensitization of meningeal afferents. Examples of experimental trials depicting the responses (post-stimulus time histogram; 0.1 sec) of two C afferents to threshold (TH) and suprathreshold (STH) mechanical stimuli (green traces) applied to the their dural receptive field during baseline recording and then at 30 minutes following CSD elicitation in control (***A***) and naproxen-treated animals (***B***). Note the lack of sensitization following naproxen treatment. Individual threshold and suprathreshold responses of sensitized and non-sensitized afferents in control and naproxen-treated animals (***C-F***). Magnitudes of CSD-related threshold (***G***) and suprathreshold (***H***) changes in mechanosensitivity depicting the inhibition of sensitization by naproxen. P values based on a Mann Whitney U-test.

### The K(ATP) channel opener Levcromakalim counteracts CSD-evoked cortical hypoperfusion and reduced tpO_2_

Naproxen may inhibit the CSD-evoked afferent mechanosensitization indirectly, by blocking the cortical hypoperfusion and reduced tissue oxygenation. However, it is also possible that the inhibition of sensitization is independent of these cortical metabolic effects, since prostanoids (elaborated by CSD) whose synthesis is blocked by naproxen, exert a direct sensitizing effect on meningeal afferents (independent of metabolic effects) (Zhang et al., 2007), and naproxen also inhibits afferent sensitization induced by a mixture of inflammatory mediators (Levy et al., 2008). We therefore asked next whether opposing the CSD-evoked cortical hypoperfusion and the associated decrease in tpO_2_ using an alternative, cyclooxygenase-independent approach, might also inhibit the afferent sensitization response. We postulated that local application of a potent cortical vasodilator could serve this purpose and tested the effects of levcromakalim, a selective ATP-sensitive potassium channel (K(ATP) opener with strong cortical vasorelaxation properties (Reid et al., 1995). In preliminary experiments, we observed that the maximum cortical vasodilatory effect of levcromakalim occurs within 15-25 min. Therefore, in order to counteract the CSD-evoked persistent hypoperfusion, we used a pretreatment approach, and started levcromakalim treatment 30 min before CSD elicitation. Using this protocol, prior to CSD induction, levcromakalim (n=28) increased basal cortical blood flow [196.10% (72.7)] [median (IQR)]; (W=231.0; p<0.0001, Figure 7A). Levcromakalim did not inhibit the elicitation of CSD (n=3, Figure 7C), which was associated with a significant cortical hyperemic response, albeit smaller in magnitude when compared to the response in control animals [105.3% (1.9)] vs [180.3% (61.6)] [median (IQR)]; (U=0.0; p<0.0001; Figure 7E). In the presence of levcromakalim, there was a decrease in cortical blood flow following CSD, but values did not attain the hypoperfusion level observed in control animals (Figure 7F; U=51.0; p<0.0001); up to 45 min post-CSD period values were not different than those observed during the pre-drug baseline period [99.56% (39.67] [median (IQR)]; (W=-8.0; p>0.99), and increased afterwards [119.1% (54.9)] [median (IQR)]; at 60 min post-CSD; (W=366.0; p<0.0001).

**Figure 7.**
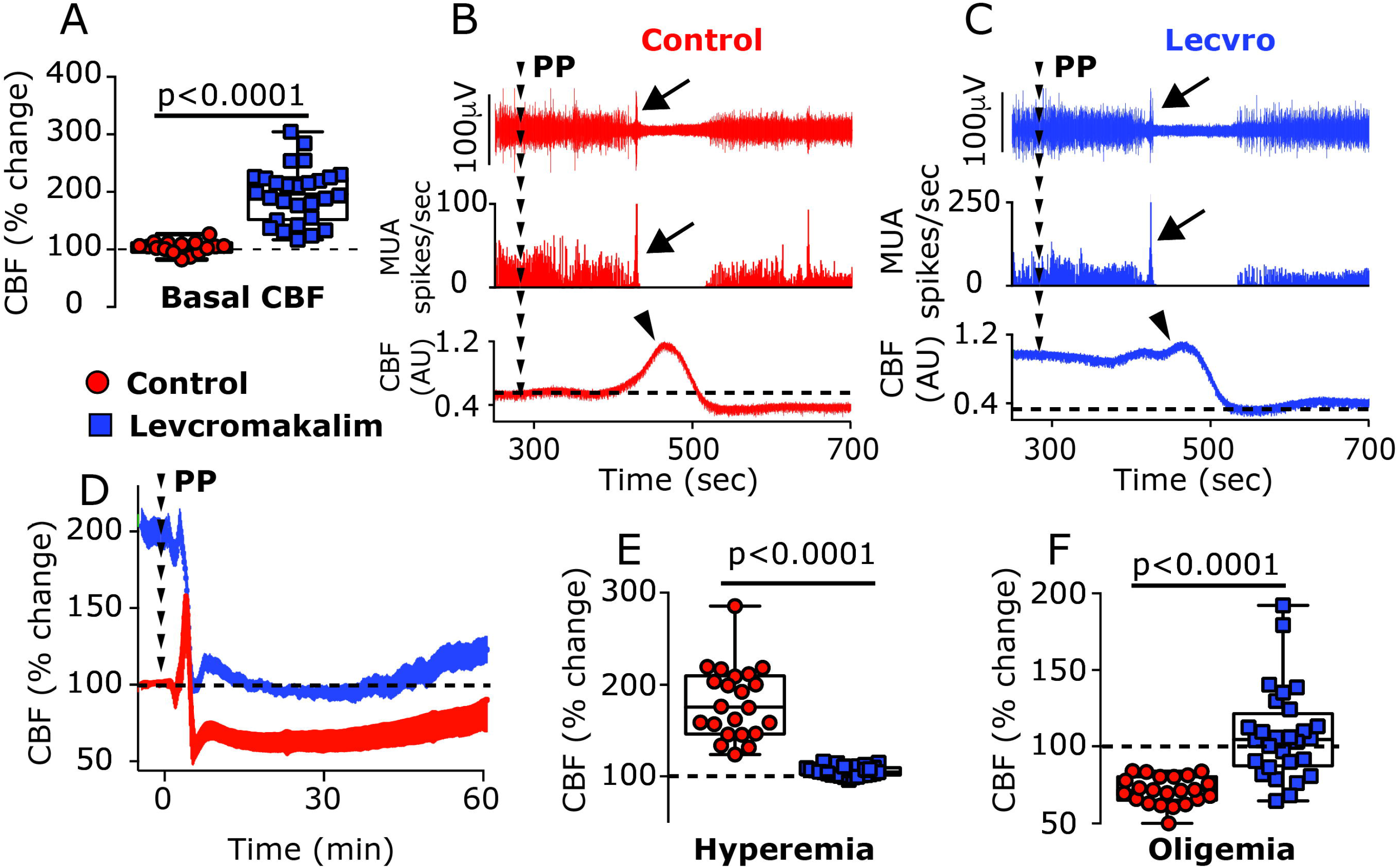
Effects of the K(ATP) channel opener levcromakalim on CSD elicitation and CSD-evoked CBF changes. **A**. Levcromakalim increased basal CBF. Examples of combined recording of multiunit recording of cortical neuronal activity (post-stimulus time histogram; 1 sec in middle panels) and CBF (bottom panels) showing a similar induction of CSD in control (***B***) and levcromakalim-treated animals (***C***). Note the initial depolarization (increased cortical activity; arrows) followed by the suppression and gradual recovery of neural activity. Also note the CSD-evoked cortical hyperemic response in the control animal and the diminished response in levcromakalim-treated animal (arrowheads). ***D***. Time-course changes of CBF in control and levcromakalim-treated animals following PP-induced CSD. Note that despite the decrease in CBF in levcromakalim-treated animals, values remained near pre-drug baseline values (denoted as 100%) unlike the marked decrease in control animals. ***E***. CBF changes in control and levcromakalim-treated animals during the hyperemia stage. ***F***. CBF changes during the hypoperfusion phase, 10-60 min following CSD. P values based on a Mann Whitney U-test.

We next tested whether levcromakalim might also affect the CSD-evoked changes in cortical tpO_2_ (n=8). Prior to CSD elicitation, levcromakalim increased basal tpO_2_ [Δ43.55 mmHg (35.3)] [median (IQR)]; (W=36.0; p=0.008; Figure 8A). In the presence of levcromakalim, CSD gave rise to a further, brief increase in tpO_2_ [peak 120.5 mmHg (56.3)] [median (IQR)]; (W=36.0; p=0.008; phase 1 in Figure 8A) with a magnitude similar to that observed in control animals [Δ18.22 mmHg (10.32)] vs [Δ14.18 mmHg (3.87)] [median (IQR)]; (U=24.0; p=0.11). While the duration of this hyperoxic phase was shorter than in control animals [70.0 sec (30.0)] vs [100 sec (50.0)] [median (IQR)]; (U=11.0; p=0.05), overall the hyperoxia phase (AUC) was not different between the two groups (U=33.0; p=0.39; Figure 8C). Following the brief hyperoxia, tpO_2_ values decreased transiently, reaching hypoxic values [18.5 mmHg (26.36)] [median (IQR)], not different than in control animals [17.66 mmHg (18.64)] [median (IQR)]; (U=32.0; p=0.35). However, when compared to control animals, levcromakalim shortened this hypoxic phase [20 sec (27.5)] vs [110 sec (70.0)] [median (IQR)]; (U=3.5; p=0.0002), yielding a reduced hypoxic response [246.6 mmHg*sec (676.0)] vs [2001.0 mmHg*sec (4883.9)] [median (IQR)]; (U=7.0; p=0.002; Figure 8D). Levcromakalim also prevented the prolonged reduction in tpO_2_ when compared to control animals (U=15.0; p=0.001; Phase 3 in Figure 8B, Figure 8E), with values not different than baseline at the 5-10 min [49.49 mmHg (43.2)] [median (IQR)]; (U=26.0; p=0.46), and 10-20 min time-points [47.16 mmHg, 44.51)] [median (IQR)]; (U=31.0; p=0.64).

**Figure 8.**
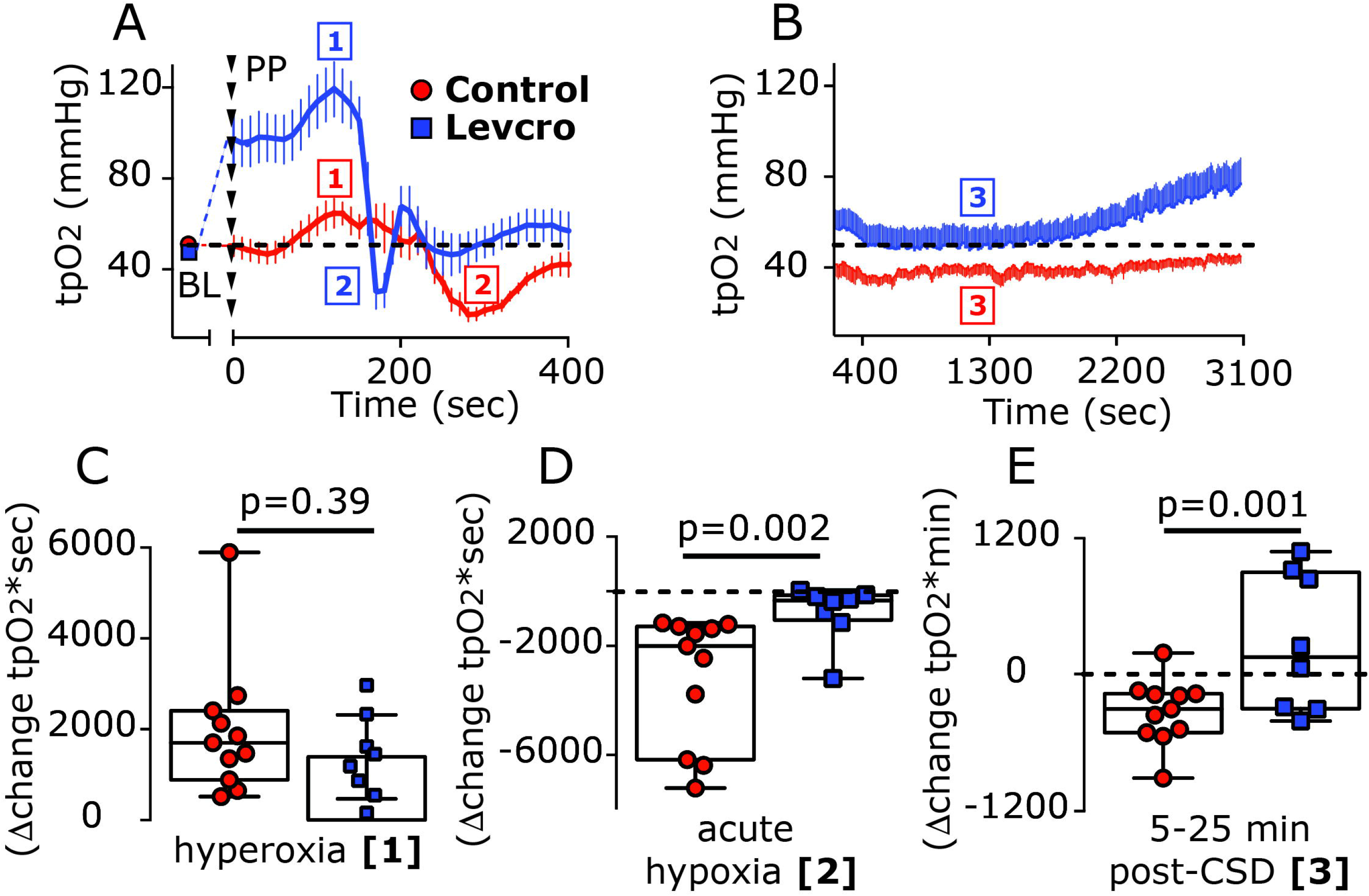
Levcromakalim inhibits CSD-evoked cortical hypoxia. Time-course changes (median and IQR) in cortical tpO_2_ values at baseline (BL) and during the acute hyperoxia and hypoxia phases in response to CSD induction with PP (phases 1 and 2 respectively) in control and levcromakalim-treated animals (***A***). Note the shorter duration of the hypoxia phase following levcromakalim treatment. The prolonged, post-CSD decrease in tpO_2_ (phase 3) in control animals and its amelioration by levcromakalim are shown in ***B***. Dashed line represents pre-drug baseline values. Differences between control and levcromakalim-treated animals in the hyperoxic response (Area under the Curve, AUC, phase 1; ***C***), acute hypoxia (AUC, phase 2; ***D***), and the prolonged decrease in tpO2 at 5-25 min post-CSD (AUC, phase 3; ***E***). P values based on a Mann Whitney U-test.

### Levcromakalim inhibits CSD-evoked activation but not sensitization of meningeal afferents

Given its ability to counteract the cortical metabolic perturbations evoked by CSD, we next determined the effect of levcromakalim treatment on the development of CSD-evoked afferent activation in 9 A-delta and 11 C afferents. In the presence of levcromakalim, baseline ongoing activity in A-delta [0.13 spikes/sec (0.31)] [median (IQR)] and C afferents [0.28 (0.89)] [median (IQR)] were not different than in control animals (U=91.0; p=0.32 and U=156.0; p=0.41 respectively). Levcromakalim treatment, however, significantly inhibited the acute afferent activation [A-delta; 0/9 vs 9/26 A-delta in the control group; (χ(1) = 4.19, p=0.04); C afferents; 1/11 vs 14/34 in the control group; (χ(1)=3.85, p=0.049); combined population 1/20 vs 23/60; (χ(1) = 7.937; p=0.005); Figure 9C]. Levcromakalim also inhibited the prolonged afferent activation [A-delta; 1/9 vs 13/26; (χ(1) = 4.21; p=0.04); C afferents; 3/11 vs 22/34; (χ(1) = 4.717; p=0.03); combined population; 4/20 vs 35/60; (χ(1)=8.82; p=0.003); Figure 9D].

**Figure 9.**
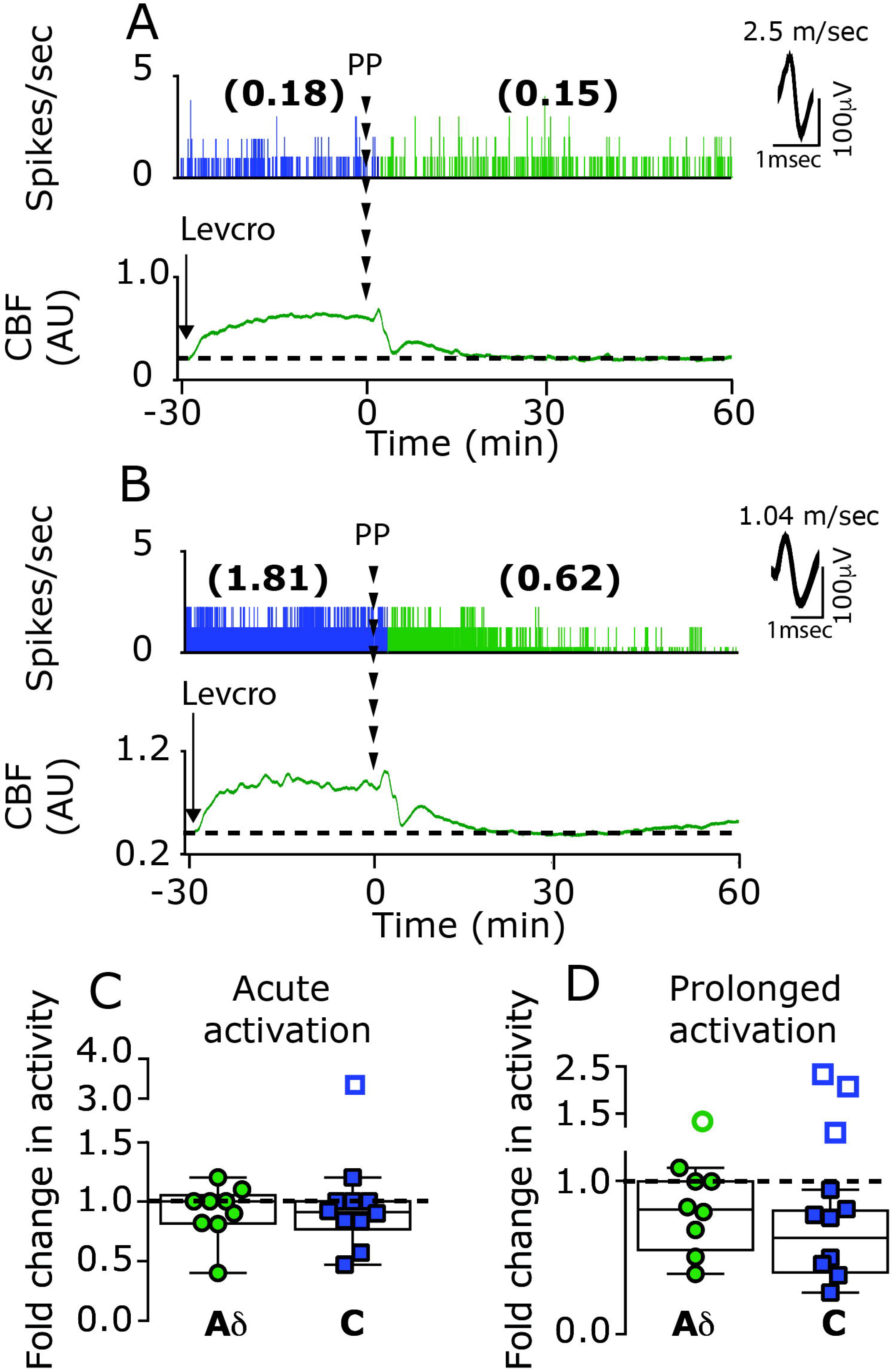
Levcromakalim inhibits CSD-evoked acute and prolonged meningeal afferent activation. Examples of single-unit recording of an A-delta afferent (***A***) and a C afferent (***B***), combined with laser Doppler flowmetry, depicting the lack of acute and prolonged afferent following the elicitation of CSD with PP in animals treated with levcromakalim. In the presence of levcromakalim, only one C afferent became acutely activated (open square outside the min/max box in ***C***.). Prolonged activation was noted only in 1 A-delta and 3 C afferents (open circle and squares outside the min/max box in ***D***.)

Finally, we tested the effect of levcromakalim on the development of CSD evoked mechanical sensitization in 7 A-delta and 10 C afferents, which had a similar number of identified dural mechanosensitive receptive fields and baseline mechanical responsiveness as the afferents tested in the control group (Table 1). Unlike its effect on the CSD-evoked afferent activation, levcromakalim treatment did not inhibit the sensitization response (see an example in Figure 10A). When compared to control experiments, levcromakalim did not reduce the relative number of afferents that were deemed sensitized when using threshold [(10/17 vs 13/23 affected afferents; χ(1) = 0.04; p=0.85)] or suprathreshold [(6/17 vs 14/23 affected afferents; χ(1) = 1.1, p=0.29)] stimuli. When sensitization was defined as enhanced responsiveness at either stimuli level, the ratio of sensitized/non-sensitized was also not different when compared to that observed in control animals [11/17 vs 17/23; (χ(1) = 0.2, p=0.66)]. When compared to control animals, levcromakalim also did not decrease the magnitude of the sensitization responses to threshold [Δ1.73 spikes/sec (3.36)] vs [Δ1.3 spikes/sec (2.18)] [median (IQR)]; (U=38.5; p=0.10; Figure 10D), or suprathreshold stimuli [1.39 fold (0.5)] vs [1.37 fold (0.54)] [median (IQR)]; (U=34.0; p=0.54; Figure 10E).

**Figure 10.**
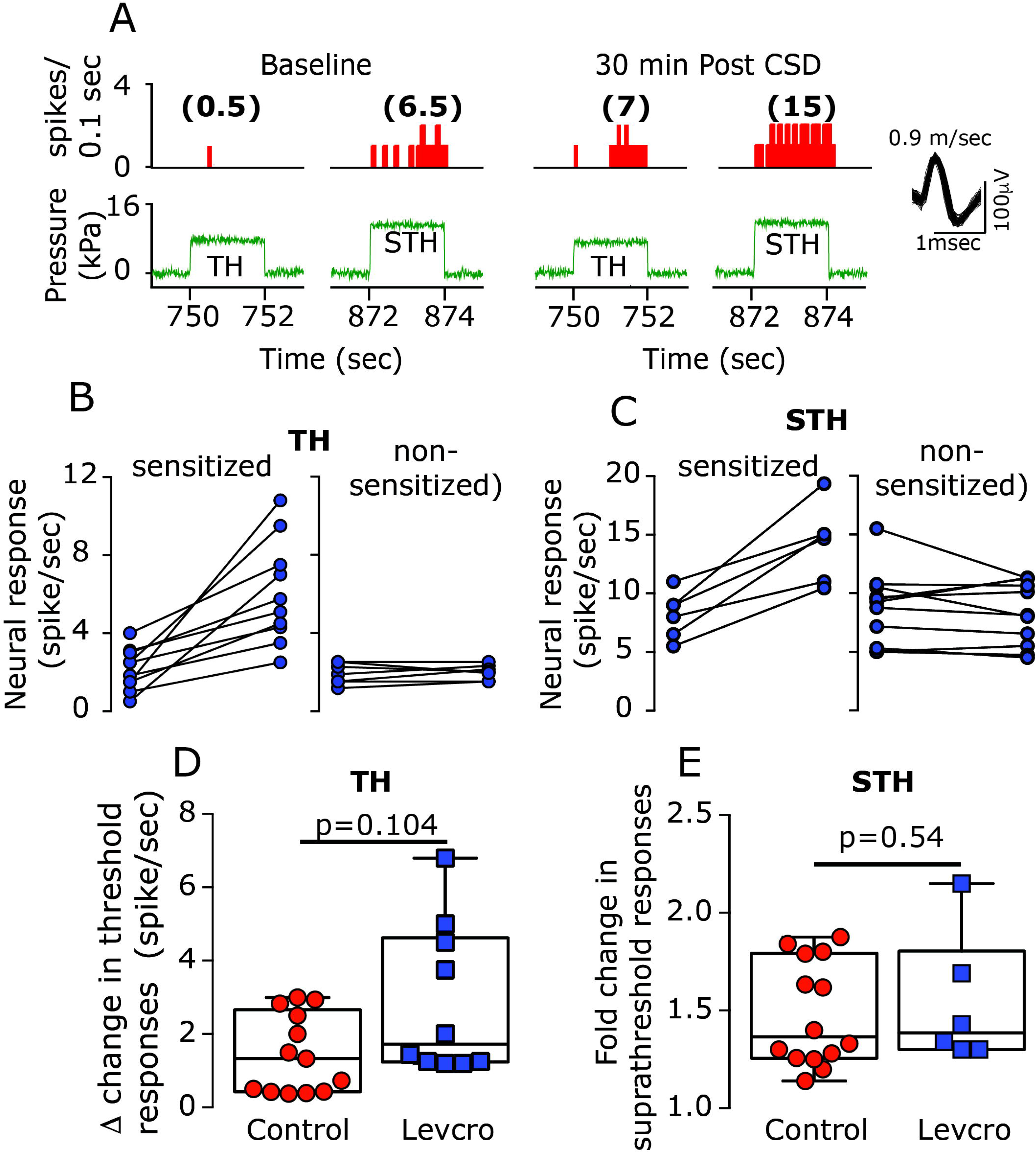
Levcromakalim does not inhibit CSD-evoked mechanical sensitization of meningeal afferents. ***A***. An example of an experimental trial depicting the CSD-evoked increase in mechanical responsiveness of a C afferent recorded from a levcromakalim-treated animal. Note that CSD-evoked sensitization at both the threshold and suprathreshold levels (post-stimulus time histogram; 0.1 sec). Individual threshold (***B***) and suprathreshold (***C***) responses of sensitized and non-sensitized afferents. Comparisons between the magnitudes of CSD-related changes in mechanosensitivity at the threshold (***D***) and suprathreshold levels (***E***) depicting similar CSD-evoked sensitization responses in control and levcromakalim animals. P values based on a Mann Whitney U-test.

## Discussion

Spreading cortical hypoperfusion has been demonstrated during the early stage of migraine with aura and likely reflects a CSD-related vascular event (Ayata and Lauritzen, 2015). And decreased cortical oxygenation has been suggested to trigger migraine headache (Schoonman et al., 2006; Arngrim et al., 2016). In our CSD rodent model, the cortical hypoperfusion and decreased tpO_2_ coincided with the emergence of the prolonged activation of meningeal afferents, suggesting a potential interaction. However, a key finding of this study was that despite exerting a powerful inhibitory effect on the protracted cortical hypoperfusion and the related decrease in oxygenation, naproxen did not ameliorate the activation of meningeal afferents in response to CSD. Our data thus point to a dissociation between these CSD-related cortical metabolic perturbations, or downstream processes, and the mechanism responsible for driving meningeal afferent activity. The ineffectiveness of naproxen was somewhat surprising in light of its ability to reduce the persistent activation of meningeal afferents following application of a mixture of inflammatory mediators to their meningeal receptive field (Levy et al., 2008) – a model of meningeal nociception (Strassman et al., 1996) and migraine pain (Oshinsky and Gomonchareonsiri, 2007; Edelmayer et al., 2009; De Felice et al., 2013). This discrepancy suggests that the prolonged activation of meningeal afferents in the wake of CSD occurs via a distinct mechanism, one that does not involve local nociceptive actions of these classical inflammatory mediators (Strassman et al., 1996) or vasoconstricting prostanoids that mediate the cortical hypoperfusion following CSD (Shibata et al., 1992; Gariepy et al., 2017).

The cortical hyperemia that occurs during the CSD wave is likely driven in part by the acute activation of meningeal afferents, and the ensuing release of vasodilatory neuropeptides such as CGRP (Busija et al., 2008). We found that naproxen also increased the initial cortical hyperemia, which likely mediated the conversion of the acute hypoxia phase to hyperoxia, but these effects did not influence the acute activation of the afferents. These data support the view that cortical vasodilation and its related increases in tissue oxygenation do not drive meningeal afferent activity (Levy and Burstein, 2011) and that during the hyperemic phase of the CSD, there is a concurrent release of opposing vasoconstricting prostanoids.

During the prolonged cortical hypoperfusion phase, CSD also gives rise to persistent vasodilatation of the middle meningeal artery (Bolay et al., 2002), a vascular response proposed to be mediated by prolonged activation of dural afferents and downstream signaling via a central parasympathetic reflex (Bolay et al., 2002). Importantly, a recent study demonstrated that this CSD-evoked meningeal vasodilatory response could be blocked by systemic administration of naproxen (Karatas et al., 2013). That naproxen did not inhibit the CSD-evoked activation of meningeal afferents together with previous data showing its inability to block parasympathetic outflow to the cranium (Akerman et al., 2012) suggests therefore that CSD-evoked dural vasodilatation may not be driven by enhanced meningeal afferent ongoing activity. Our present results thus call into question the use of dural artery vasodilation as a surrogate of increased meningeal afferent activity and the activation of the migraine pain pathway in preclinical studies.

We found that counteracting the protracted cortical hypoperfusion and prolonged decrease in tpO_2_, by preemptively increasing cortical blood flow using levcromakalim, was associated with inhibition of the acute and prolonged afferent activation phases. We propose however that the inhibitory effect of levcromakalim on these afferent responses did not involve the normalization of cortical blood flow or tpO_2_ given that naproxen, which abrogated the cortical metabolic perturbations following CSD, failed to inhibit the afferent activation. We cannot exclude the possibility that the cortical vasodilation and increased tpO_2_ induced by levcromakalim, prior to the onset of CSD, contributed somehow to the blockade of the post-CSD afferent activation. However, the finding that levcromakalim did not affect the afferents’ basal ongoing activity suggests that the vascular effects of levcromakalim were unlikely to mediate to the inhibition of the CSD-evoked afferent activation.

Cortical neurons also express K(ATP) channels (Karschin et al., 1997) and their prolonged activation by levcromakalim could have resulted in diminished cortical excitability (Soundarapandian et al., 2007), potentially affecting the elicitation of CSD. However, levcromakalim did not inhibit the induction of CSD, and particularly not the initial strong depolarization prior to the prolonged suppression of cortical neural activity. K(ATP) activation in cortical neurons could, nonetheless, reduce the release of pro-nociceptive transmitters, such as glutamate (Zhao et al., 2013), that may contribute to the acute and potentially the prolonged activation of meningeal afferents. Cortical astrocytes also express K(ATP) channels (Thomzig et al., 2001) and their activation could potentially decrease the high extracellular K^+^ level that is attained during the passage of the CSD wave (Kraig and Nicholson, 1978) by enhancing K^+^ “spatial buffering” (Kofuji and Newman, 2004). In response to CSD, elaborated K^+^ in the cortical extracellular space has been suggested to diffuse into the leptomeninges and promote the acute activation of meningeal afferents and potentially also their prolonged activation, at least in part (Pietrobon and Moskowitz, 2012; Zhao and Levy, 2018). Activation of astrocytic K(ATP) channels, leading to decreased parenchymal K^+^ thus could have potentially inhibited the CSD-evoked afferent activation. Our recent finding that K^+^-evoked acute activation of meningeal afferents was not associated with mechanical sensitization (Zhao and Levy, 2018) is also in agreement with the observation that levcromakalim did not inhibit the CSD-evoked mechanical sensitization response. Finally, activation of K(ATP) channels expressed at the peripheral nerve endings of meningeal afferents could have also contributed to the inhibition of the afferent activation by reducing their overall excitability (Chi et al., 2007). However, the finding that levcromakalim did not reduce basal ongoing discharge or inhibit the mechanical sensitization argues against such a mechanism.

We found that naproxen distinctly inhibited the CSD-evoked mechanical sensitization of meningeal afferents. That levcromakalim treatment greatly reduced the acute hypoxia phase, and normalized the prolonged cortical hypoperfusion and decreased tpO_2_, but did not inhibit the enhanced mechanical responsiveness suggests however that the mechanism underlying the mechanical sensitization following CSD is unrelated to the changes in cortical perfusion and tpO_2_. It should be noted that while levcromakalim treatment normalized cortical blood flow and tpO_2_ (to values not different than those observed before CSD), it did not inhibit the process that led to their decreases following CSD. We therefore propose that the mediators that contribute to these metabolic changes (likely cyclooxygenase-derived; see below) are responsible for the afferent sensitization rather than the cortical hypoperfusion and abnormally reduced tpO_2_ *per se.* It is possible that the acute hypoxic phase that occurred in levcromakalim-treated animals (albeit for only a fraction of the time noted in control animals) may have contributed somehow to the afferent sensitization process. However, we also observed brief cortical hypoxia in some naproxen-treated animals, further suggesting a dissociation between the acute hypoxic phase and the afferent changes.

The discrepancy between the effects of naproxen and levcromakalim on the CSD-evoked afferent activation and mechanical sensitization responses supports the view that these two migraine-related nociceptive responses are mediated by distinct mechanisms (Levy and Strassman, 2004; Zhang et al., 2010b; Zhang et al., 2013; Zhao and Levy, 2016). Naproxen’s site of action and the nature of the cyclooxygenase-derived factors that distinctly promote the sensitization of meningeal afferents following CSD require further studies. Release of cyclooxygenase-derived sensitizing mediators from cortical neurons and glial limitans astrocytes that interface with the meninges (Xu et al., 2004) could contribute to the afferent sensitization. The mode of cortex to meninges transport of cyclooxygenase-derived prostanoids could involve diffusion, or bulk flow via the glymphatic system (Iliff et al., 2012; Schain et al., 2017). The transport of sensitizing factors from the parenchymal interstitial space into the subarachnoid CSF could influence primarily leptomeningeal afferents and meningeal afferents that terminate within arachnoid granulations (von During and Andres, 1991; Fricke et al., 1997). Additional flow of sensitizing mediators from the subarachnoid CSF into the dural lymphatics system (Raper et al., 2016; Louveau et al., 2017) could affect the responses of dural afferents. Naproxen treatment could have also inhibited the CSD-evoked meningeal afferent sensitization by targeting constitutively expressed meningeal cyclooxygenase in meningeal vascular and immune cells (Zhang et al., 2009) and the ensuing release of sensitizing prostanoids (Zhang et al., 2010b).

In summary, our findings point to a dissociation between the CSD-evoked cortical metabolic perturbations and the prolonged activation and mechanical sensitization of meningeal afferents. The findings that K(ATP) activation and cyclooxygenase inhibition have distinct effects on the CSD-evoked activation and sensitization of meningeal afferents may explain the ineffectiveness of cyclooxygenase inhibitors in many cases of migraine (Suthisisang et al., 2010). A combination therapy, which includes a cyclooxygenase inhibitor and a K(ATP) channel opener, may be explored for treating the headache in migraine with aura.

## Acknowledgments

We thank Dr. A. M. Strassman for helpful comments on the manuscript. The study was supported by NIH grants NS086830, NS078263, NS101405 to DL.

